# Whales in Space: Experiencing Aquatic Animals in Their Natural Place with the Hydroambiphone

**DOI:** 10.1101/2023.12.27.573441

**Authors:** James P. Crutchfield, David D. Dunn, Alexandra M. Jurgens

**Affiliations:** Complexity Sciences Center, Physics and Astronomy Department, University of California, Davis, California 95616; Art & Science Laboratory, 613 C Street, Davis, California 95616; Inria Centre, University of Bordeaux, France

**Keywords:** marine mammals, humpback whales, undersea vocalizations, ambisonics, hydrophone array, spatial sound

## Abstract

Recording the undersea three-dimensional bioacoustic sound field in real-time promises major benefits to marine behavior studies. We describe a novel hydrophone array—the hydroambiphone (HAP)—that adapts ambisonic spatial-audio theory to sound propagation in ocean waters to realize many of these benefits through spatial localization and acoustic immersion. Deploying it to monitor the humpback whales (*Megaptera novaeangliae*) of southeast Alaska demonstrates that HAP recording provides a qualitatively-improved experience of their undersea behaviors; revealing, for example, new aspects of social coordination during bubble-net feeding. On the practical side, spatialized hydrophone recording greatly reduces post-field analytical and computational challenges—such as the “cocktail party problem” of distinguishing single sources in a complicated and crowded auditory environment—that are common to field recordings. On the scientific side, comparing the HAP’s capabilities to single-hydrophone and nonspatialized recordings yields new insights into the spatial information that allows animals to thrive in complex acoustic environments. Spatialized bioacoustics markedly improves access to the humpbacks’ undersea acoustic environment and expands our appreciation of their rich vocal lives.

## 1. INTRODUCTION

Marine mammals spend the bulk of their active lives submerged beneath the sea surface. Given the relatively poor propagation of light compared to sound in the ocean depths, the world of these animals is primarily acoustic. These factors greatly complicate relying solely on surface observations to address the full diversity of their behaviors. Fortunately, in the last decade or so scientists demonstrated the substantial benefit of undersea, comprehensive tracking with, for example, skillful attachment of digital devices that monitor animal behavior via sensors that record video, sound, location, depth, pressure, temperature, and the like [1]. The following describes complementary benefits that come from recording the underwater three-dimensional bioacoustic sound field in real-time.

### A. Whale Bioacoustics

Sound propagation in water differs from that in air: sound travels five times faster in water than in air, acoustic waves in water propagate with much less dissipation, and different frequencies travel at different speeds. (See Table I.) These phenomena make undersea sound markedly more complex to analyze, understand, and harness. They complicate directly monitoring and interpreting sound in the ocean. That said, these properties also mean there is additional information available in ocean acoustic waves to be harnessed for environmental sensing and for communication. (See App. A.)

To begin to address these challenges, we applied spatial bioacoustics to monitor humpback whales (*Megaptera novaeangliae*) of southeast Alaska, demonstrating that it markedly improves understanding their undersea behaviors. As one example, the following describes how acoustic spatialization revealed previously unreported aspects of social coordination during bubble-net feeding [2].

Cetaceans exhibit compelling evidence for advanced intentional behaviors and conscious awareness through their raw intelligence, song generation [3, 4] and sharing [5, 6], communication and interactions with their own and other species [7, 8], and empathy (concern for others’ well-being) [9]. Humpback whales, in addition, are known to be very vocal and social [10].

Evolving over a time span ten times that of humans, cetaceans developed tools (socially-coordinated bubble-net feeding by humpbacks) and region- (and possibly hemisphere-) spanning ocean-acoustic communication networks [11]. Over the last half century humpback whales, in particular, became known for their active vocalizations. These fall into two categories: One comprised of extended *songs* (minutes to hours), emitted predominantly by males; the other *social calls*, short vocalizations (lasting seconds) that occur in animal interactions and are produced by both males and females [3].

**TABLE I:** Sound propagation differences in air and water: (i) Medium density, (ii) medium bulk modulus, (iii) sound velocity, and (iv) sound wavelengths at two different frequencies. Unlisted, but important is sound dispersion: the range of frequency-dependent velocities is markedly large in water. ^*^Standard atmospheric conditions (0 C or 32 F, at sea level). ^**^Standard atmospheric pressure (1 atm.). ^***^ At sea surface. ^****^ Room temp (20 C, 68 F).

| Medium | Density<br>(kg/m<sup>3</sup>) | Bulk Modulus<br>(Pa) | Sound Velocity<br>(m/s) **** | Wavelength<br>100 Hz (m) **** | Wavelength<br>1000 Hz (m) **** |
| --- | --- | --- | --- | --- | --- |
| Air | 1.225 * | $1.42 \times 10^5$ | 343 | 3.43 | 0.343 |
| Fresh Water | 1000 ** | $2.15 \times 10^9$ | 1482 | 14.82 | 1.482 |
| Seawater | 1025 *** | $2.29 \times 10^9$ | 1500 | 15 | 1.5 |

Song function is still largely a mystery. Since songs are predominantly produced by males, historically they have been interpreted as facilitating mate choice and so playing a role in sexual selection [12–14]. However, more recent results suggest that humpback songs are not so much about reproductive fitness, rather they may “reveal the precise locations and movements of singers from long distances and may enhance the effectiveness of [acoustic] units as sonar signals” [15]. That is, vocalization is important to navigation which is central to the very long seasonal migrations (1000s km) of humpback whales.

These debates highlight the need to discover patterns and the information contained in animal vocalizations and to place these tasks at the center of monitoring and interpreting cetacean behavior. This, in turn, calls for more study and new acoustic instrumentation that reveals spatial aspects of whale vocalizations. And, this suggests exploring new concepts of patterns and structure in data as developed in the mathematical theory of causal statistical inference [16–18] and algorithmic approaches from modern machine learning and AI (ML/AI) [19].

Both the foundational theory and ML/AI methods require substantial datasets and in the latter case typically require labeling by human experts. We show that new kinds of data from new kinds of instrumentation can be equally or more important to interpreting acoustic data than exploiting massive data, algorithm advances, and huge computational resources.

The principal reason to pursue innovations in instrumentation is that acoustic spatialization disambiguates the locations of animal vocalizations, markedly improving access to the humpbacks’ undersea acoustic environment and expanding our appreciation of their rich vocal lives. In addition, spatialized hydrophone recording greatly reduces the post-field analytical and computational challenges when confronted by many vocalizers and multiple additional sound sources, as commonly occurs in field recordings. In contrast to these well-known difficulties, it is important to keep in mind that human observers and apparently marine mammals are not prohibitively challenged by multiple-source ambiguity—the so-called “cocktail party problem”.

The following introduces a novel hydrophone array— the *hydroambiphone*—that adapts ambisonic spatial-audio theory to sound propagation in ocean waters to largely alleviate such problems, while providing (i) clues to the kinds of information that allow animals to thrive in complex acoustic environments and (ii) markedly more representative experiences of their acoustic world.

### B. Undersea Ambisonics

To explore the complexities of the undersea acoustic environment and circumvent these problems, we adapted the ambisonic theory of spatial acoustics [20, 21]. The theory formalizes the representation of an ambient sound field systematically in terms of three-dimensional spherical harmonics—the solutions of the wave equation that describes sound propagation in a medium. The theory gives an exact representation of a sound field centered at a given point in space in terms of a systematic approximation at an infinite number of “orders”. In practice one can only go to finite-order approximation. The higher the order, though, the better the spatial resolution of the approximation, but the number of required transducers grows rapidly with order. For this reason, real applications generally use low-order approximations. Ambisonic theory applies to both recording and playback of three-dimensional spatial sound fields.

In-air ambisonic recording arrays use directional microphones—cardioid or super-cardioid, for example. They are placed and oriented to completely cover a sound field with non-overlapping regions. This is not possible in water, since the available transducers—hydrophones— are omnidirectional. To address this our implementation uses a first-order ambisonic approximation consisting of four omnidirectional hydrophones mounted on the surface of a 12-inch diameter hollow sphere. Given the corrosive nature of seawater the sphere was stainless steel. It also provided a high specific acoustic impedance to isolate hydrophones from acoustic waves propagating from directions opposite each hydrophone. Thus, the spherical shape provides a kind of “acoustic shadowing” that improved the directivity of each hydrophone, whose individual sensitivity is otherwise omnidirectional, as noted.

The following demonstrates how to adapt ambisonics to the ocean acoustic environment and proves out the HAP as an effective marine bioacoustic instrument. The purpose in this is two-fold: (i) introduce the HAP and describe its deployment and (ii) recount several novel scientific results from our August 2023 voyage that established its use as a viable marine science instrument.

To these ends, we briefly layout HAP design (Sec. II), calibration and performance (Sec. III), spatial audio signal processing (App. D), and listening (App. E). We do cite, however, companion technical descriptions that go into considerably more detail. The main focus here are the scientific results that came from using the HAP’s spatial audio: (i) access to a new immersive experience of the humpback acoustic Umwelt, (ii) novel acoustic coordination during bubble-net feeding, (iii) dynamics of group breaching, (iv) HAP high acoustic sensitivity, (v) undersea infrasonics, and (vi) undersea noise in the Inside Passage.

### C. Overview

The following recounts the results of deploying the HAP during a 300 mile voyage along the Inside Passage of southeast Alaska 18 August to 2 September 2023, transiting from Juneau to Ketchikan aboard the M/Y Blue Pearl (Don and Denise Bermant, owner/operators).

First, we review ambisonic spatial audio theory and then move on to describe how this is adapted to the design of a hydrophone array that records the undersea three-dimensional sound field. We comment on several design aspects, its testing and calibration, and required digital audio recording and playback for listening. We summarize the HAP’s performance in Sec. III B.

We then review a selection of new undersea bioacoustic phenomena, most related to humpback whale behaviors and vocalizations, but also recount several notable purely ocean-acoustic observations. This includes a brief discussion of the present challenges. We end by drawing out a number of conclusions that range from emphasizing the novelty of the whale bioacoustics identified to suggesting future prospects for greatly improved HAP systems and the possible innovations in marine bioacoustics.

## II. CAPTURING SPATIAL BIOACOUSTICS

A sound field is a three-dimensional organization of acoustic energy sustained by oscillatory motions of a medium, such as air or water. When a sound source activates, the field agitation propagates in all directions away from the source in spatiotemporal patterns governed by the wave equation: 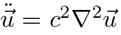, where *c* is the speed of sound and 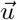 (*x, y, z*) is local state of the medium. The result is that monitoring the instantaneous state of a time-dependent sound field requires measuring and then recording a (vector valued) function 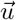 throughout a three-dimensional volume.

When we listen to an audio recording of a musical performance, our ears are presented with a sound field that is generated by a particular spatial configuration of voltage-to-pressure transducers (loudspeakers) that are driven by signals picked up by artfully-placed individual sound-to-voltage transducers (microphones). It is one of the sound recording engineer’s primary responsibilities to determine the types of microphone and their placement— a contact small-diaphragm condenser mic near sound hole of the acoustic guitar, a dynamic mic near the singer’s mouth, a unidirectional cardioid close to and facing the drums—to best capture and then reproduce the musical experience.

Unfortunately, the selected mic placements anchor the recorded sound signals to those specific locations. Once recorded, those locations cannot be changed. Post-recording, there is little variation available to acoustically explore, since so much of the original ambient 3D sound field 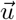 information is lost in a collection of spatially-local recordings. Moreover, careful and extensive mic placements are typically not feasible nor is the resulting sound reproduction adequate for natural sound fields.

### A. Ambisonics

An alternative way to capture a 3D sound field is afforded by adapting ambisonic (spatial audio) theory [20, 21]. Its benefits include (i) listener-centric representation of the sound field (rather than a set of mic-centered signals), (ii) flexible post-recording signal processing, such as panning, zoom, rotation, and beam forming, (iii) flexible post-recording encodings to an unrestricted array of playback systems—from monophonic and stereo to modern full immersion systems with dozens or even hundreds of loudspeakers. As we will show, these benefits are key to processing and interpreting vocalizations of animals, especially those that move in three dimensions.

Ambisonics does this by expanding the sound field in spherical harmonics centered around the listening point (or *sweet spot*). These spherical harmonics are the solutions to the 3D wave equation above and give a mathematically-consistent and systematic approximation basis for representing the ambient sound field up to a given *ambisonic order*.

### B. Hydroambiphone

Ambisonic recording produces a time series of measurements each of which is vector of acoustic signals. The dimension of the vector grows quadratically with the approximation order. Thus, implementing an ambisonic transducer entails practical trade-offs between, for example, the expense and cabling of multiple transducers, large data storage requirements, and computational complexity and computing resources required to process long-duration vector signals—in real-time and off-line. Balancing these, we selected a first-order ambisonic (FOA) array consisting of four hydrophones mounted on the surface of 12 inch-diameter hollow sphere. To properly “shadow” each hydrophone from sounds arriving from opposite directions (and so improve directionality) the sphere was 2mm thick stainless steel, which has a high specific acoustic impedance. (See Apps. B and C and Fig. 1.)

**FIG. 1:**
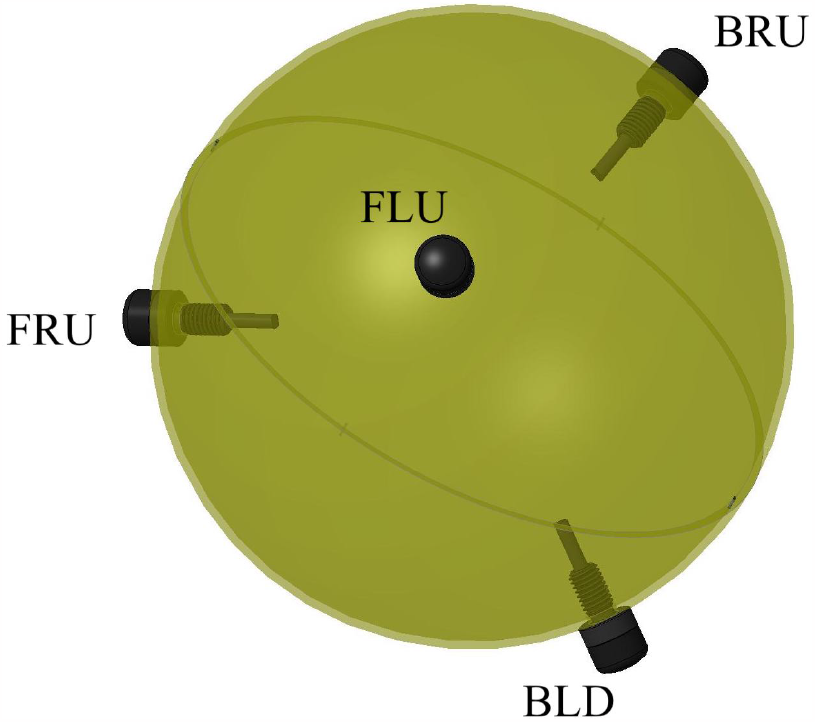
Hydroambiphone with hydrophone placements on the surface of a hollow sphere: (i) Front-Left-Up (FLU) (ii) Front-Right-Down (FRD), (iii) Back-Left-Down (BLD), and (iv) Back-Right-Up (BLU).

### C. Ambisonic Recording and Signal Processing

The HAP acoustic signals were captured by a multi-channel digital audio recorder (Zoom F8n Pro) to high-capacity, high-speed SD cards. Each of the HAP’s four channels was sampled at 24 bit resolution at a rate of kHz in the recorder’s Ambisonic mode. After a day’s collection of recording sessions, the audio data was transferred to a laptop and also to an additional backup disk. Figure 4 shows the complete first-order Ambisonic recording and observation system.

The spatial audio processing chain consisted of a specific series of stages:

1. Cleaning each HAP Ambisonic A mono channel to attenuate interference from vessel operating subsystems, such as sonar, and also from surface wave noise.
2. Converting the Ambisonic A Format vector time series to Ambisonic B (Fuma) format, accounting for the physical properties of sound in water and the HAP’s design.
3. Beamforming the B format vector signal to increase the directionality of the encoded ambisonic signals.
4. Encoding the beamformed Fuma signal to one or another playback configuration. These included stereo, binaural, a custom 6.1 Hexagonal F encoding, and several Dolby Atmos configurations ranging from a half-dozen to two dozen speakers.

For details see App. D.

The listening experience of spatialized audio depends strongly on the playback system. Available audio formats and loudspeaker configurations offer very different degrees of spatialization and acoustic immersion. Headphone listening via the binaural encodings is convenient and captures a notable amount of the spatial audio, given that it is being heard over two-channel stereo headphones. In contrast, Hexagonal F surround presents a richer sound environment with much improved source localization. Our custom-built Hexagonal F 6.1 surround playback systems consist of a subwoofer and six co-planar satellite speakers (located at head level and centered around the sweet spot where the listener sits). Listening via the Hexagonal F system is necessary to discriminate sources in complex acoustic scenes and/or with low-level or distant acoustic sources.

## III. HAP PERFORMANCE

Our main message here is that the HAP worked to capture the undersea 3D acoustic sound field. In point of fact, its performance greatly exceeded our original predictions. That said, the field deployments also revealed a number of challenges and so opportunities for improvement.

Before recounting the scientific results, we will describe the HAP’s initial field deployment on our August 2023 voyage and mention several practical issues for effective use. Second, we will then describe the HAP’s basic and spatial acoustic performance, including calibration test and a number of experimental confounds.

### A. Field Deployment

Our field deployments of the HAP occurred over two weeks (Summer-Fall 2023) on a 300 mile voyage through the Inside Passage of southeast Alaska, transiting from Juneau to Ketchikan aboard the M/Y Blue Pearl, a 65’ Fleming motor yacht. Of note for supporting our field tests the vessel provided 115 V AC power for the recording and playback equipment and device battery charging and Starlink marine uplink to the Internet with 10 Mbps upload, 20-50 Mbps download, and very low latency.

The HAP was supported by a stainless steel cable that relieved stress from the four hydrophone signal cables. Given the 60’ long cables the HAP was deployed from 15’ - 50’ depths. It was particularly important to carefully track the vessel’s position and orientation with respect to current, wind, drift, and surface waves. Without monitoring these, the strong Alaskan currents would sweep the HAP under the vessel and wave slap against the vessel hull could become a distracting source of acoustic noise.

### B. Performance and Acoustic Confounds

Due to the focus here on the HAP’s practical use as an instrument for marine animal behavior, we briefly outline its performance. Detailed quantitative performance measurements will be reported elsewhere.

As far as detecting vocalizations from aquatic animals, the relevant aspects of the HAP’s performance include: (i) high sensitivity, (ii) long distance detection of sound sources (including marine mammals and human-generated sounds), and (iii) good directivity resolution.

The high sensitivity manifested itself by the HAP’s ability to detect extremely low-level sound sources— including both sources that originated close to the vessel, such as wave noise from the vessel’s hull, and sources, such as engine noise, from other vessels over a dozen miles away (well beyond the horizon and simply not visible).

Initially, the HAP’s high sensitivity seemed detrimental to the recordings as it added many nonbiological sources not of direct interest. For example, one persistent problem in the Inside Passage was the (even very distant) transit of cruise liners. Their noise amplitudes at the HAP were so large that we typically aborted recording sessions as the animal vocalizations could be entirely masked and the incoming signals were painful for human listeners. Subsequently, however, we were able to remove and attenuate these noise sources during post-field processing largely due to the Ambisonic benefits noted above.

An important parameter of the HAP was its directional sensitivity: how close (in solid angle) can independent sources be and still be distinguished. Pre-field tests in our lab tanks indicated being able to distinguish sources at 90 or more degrees of solid angle—octants on a sphere centered at the HAP. However, field tests with the vessel’s tender circulating at various radii indicated 45 degrees of solid angle distinguishability.

## IV. UNDERSEA ACOUSTICS REVEALED BY THE HAP

On our two week voyage and over dozens of recording sessions, the HAP proved itself to be a remarkably sensitive, flexible, and easy-to use instrument for capturing the undersea three-dimensional acoustic world. This led to a number of insights and discoveries, ranging from diverse animal behaviors and vocalizations to providing spectacular undersea immersive audio listening. These included: a wholly new appreciation of the whales’ acoustic *Umwelt*; humpback pairs harmonizing their social calls during collective bubble-net feeding; the HAP’s high acoustic sensitivity over long baselines; the dynamic interplay of humpback group breaching; extensive recordings of undersea noise sources; and the detection of undersea infrasonics (sound frequencies under 20-30 Hz). We briefly outline each, leaving fuller accounts to follow-up companion articles.

### A. Intangibles made Present

As observers listening to the HAP stereo signal in real-time, were daily surprised by the high degree of acoustic activity. Due to this, the voyage resulted in over 70 HAP recordings (from 10s of minutes to an hour in length each), in toto representing many dozens of hours of observation and recording. By removing silent sections in post-field processing, we compiled the collection of recordings into a single immersive audio file two hours and forty-seven minutes long that highlights the huge diversity of humpback vocalizations.

Overall, the compilation provides a thorough-going immersion into the acoustic environment experienced by southeast Alaskan humpbacks. While listening on-board in real-time was surprising in a number of ways, the compilation recording, being a concentrated presentation of selected examples, led to even more insights. We were regularly surprised at the number of animals around us. This activity was far in excess of the numbers we expected from our visual surface observations. Another general observation was the shear diversity of humpback social calls and the regularity of apparent acoustic communication. It became clear that their vocal activity was quite high. This stands in contrast to the oft-expressed belief that, due to their preoccupation with feeding, during the summer months in southeast Alaska humpbacks are markedly less vocal than during the winter mating season at low latitudes in Hawaii.

The overall take-away message is a holistically and deeply enriched experience of the humpbacks’ spatial acoustic world–their *Umwelt* [27]—that is difficult to express in the written words. To demonstrate, we are making public a short 20 minute excerpt from the compilation that presents several highlights, available [World Wide Whale](http://demon.csc.ucdavis.edu/~chaos/share/www/WorldWideWhale/World_Wide_Whale.html).

### B. Humpback Harmonies

Spatializing the social calls used by humpbacks during bubble-net feeding revealed previously-unreported vocal coordination of different individuals. Specifically, by listening to the immersive audio encoding (binaural but especially the Hexagonal F surround system) and closely inspecting spectrograms of the section of bubble-net feeding, revealed that a second humpback joins the main feeding call just before the end of the coordinated feeding activity.

The spectrogram analysis used the unaltered HAP Ambisonic A recordings corresponding to original channel numbers. See Fig. 2 for the spectrogram of HAP channel 1. The spectrogram makes it clear when the second whale vocally differentiates: the two animals’ harmonics separate and go in two opposite directions at the very end. Moreover, listening to the spatialized recording makes it clear that the source of the vocalizations consists of two distinct animals at different locations.

**FIG. 2:**
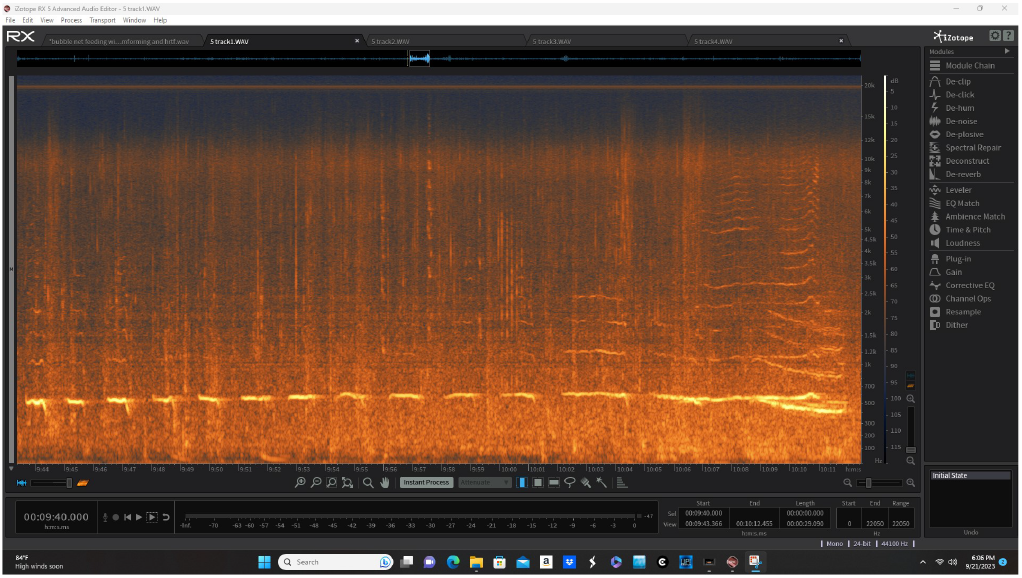
Spectrogram of a group bubble-net feeding session in which a second whale joins the existing main coordination feeding call vocalized by a different animal. Spectrogram calculated from the unaltered HAP Ambisonic A recording on HAP channel 1 only. Note where the second whale joins and how the harmonics separate and go in two directions at the very end: one up in frequency and the other down. (Reproduced here from Ref. [2] with permission.)

In addition, the recording reveals that initially, before the frequency separation, the two animal’s vocalizations start off synchronized at the same frequency. Only then does one animal shift up in frequency (as is typical in bubble-net coordination calls from lone individuals) while the other shifts down. It also clear that the phase and amplitude differences between the animals’ calls lie mostly in the mid- and high-frequency ranges, as expected. One concludes that as a vocal phenomenon these vocal coordinations are the functional equivalent of human singers harmonizing.

We recorded four such harmonizing events. Fuller description, recording data, and analysis are presented in Ref. [2]. The latter also further explores the benefit of the HAP’s spatialization that reveals the two animals are clearly vocalizing from two different locations.

### C. High Acoustic Sensitivity

The HAP revealed itself to be surprisingly sensitive, for close and very distance sound sources. For example, on board listeners often heard cruise liners and small fishing boats (their outboard motors in particular), long before they could be seen above the ocean horizon. This places these sound sources more than about 8 miles away from the vessel.

The high sensitivity at first seemed a burden, obscuring and even totally masking sounds of interest. It certainly was while listening on board. While in those situations the HAP’s high sensitivity seemed a detriment, our ultimate view is that the high sensitivity facilitates the HAP’s (and our) probing complex aspects of the 3D acoustic ocean environment in constructive ways—ways that reveal new phenomena. Specifically, when we returned post-field to the lab we developed the spatial audio processing chain described in App. D to largely remove or attenuate many of the “extraneous” sound elements that the HAP’s sensitivity introduced.

### D. Discussion

Other follow-on companion works will report on additional notable results from the HAP. These will include a spatial analysis of long periods of group breaching. From these one can estimate aspects of their configuration and even the local bathemetry. This then raises the question in a newly quantitative way of the function of breaching. Extracting spatiotemporal information from the HAP recordings in terms of contingency between breaches will allow one to probe if breaching supports communication. Another notable observation, one that at present is not explained and is counterintuitive, is repeated recording of undersea infrasonics that were highly directional. The directionality goes against conventional understanding of sound propagation since low frequencies are not associated with having a direction. Finally, over the voyage we came into contact with many natural and anthropogenic sound sources. In this way, the collection of HAP recordings can give insight into a number of undersea noise sources. We believe that the spatial aspects of these noise sources have much to contribute to our understanding of noise in the undersea environment and to monitoring how they affect the animals there.

### E. Related Work

Several independent efforts have attempted to develop devices for undersea spatial audio recording over the last decades. These range from developing directional hydrophones to hydrophone arrays. For example, Sonotronics offers their Model DH-5 Directional Hydrophone. More relevant is an implementation for stereophonic underwater sound recordings disclosed in [US Patent #8,509,034](https://patents.justia.com/patent/8509034). Finally, open source software has been developed for real-time acoustic detection and localization of cetaceans [22].

The most similar prior effort, though, is found in Refs. [23–26]. These implement an open-frame four-hydrophone array in a tetrahedral configuration—that is, a first-order ambisonic approximation. Unfortunately, the spatial localization was not strong, which appears largely due to the use of an open frame mounting for the hydrophones.

To the best of our knowledge our reports here are the first successful undersea spatial audio recording and the first used to successfully study cetacean behavior.

## V. CONCLUSION

Environmental and behavioral marine science and technology have changed immeasurably since the early days of the first appreciation of how sound propagates in the ocean and of who and what are producing those sounds. Certainly, hydrophones and ancillary signal processing have advanced substantially. These improvements promise to greatly enrich our understanding of the undersea world and its inhabitants.

We introduced a modest but accessible implementation of undersea spatial acoustics that uses inexpensive commercial-off-the-shelf (COTS) hardware and open-source software. The net functionality of the recording system allowed for real-time spatialization of aquatic animal vocalizations. We believe the HAP is a new tool for marine biology that promises to greatly expand the human appreciation of the three-dimensional acoustic world of marine animals.

Our reports of the successful proof-of-concept deployment also suggests substantial improvements. So, one can look forward to future implementations that provide increasingly higher quality and improved sensitivity and higher resolution localized spatialized sound, all available in real-time.

The operant question now is how to actively shape our future understanding of whale communication in the wild. The results suggest a coming era of citizen marine social science. The hydroambiphone is straightforward to construct at moderate cost and so gives a practical path to a new era of listening to the voices of the deep.

## ACKNOWLEDGMENTS

The authors are deeply indebted to Don and Denise Bermant for their generous tour of the Inside Passage, southeast Alaska, on the M/Y Blue Pearl (Vancouver, British Columbia) and their skillful navigation, marine insights, and patience, as well as camaraderie. All were key to the voyage’s success. They also thank Anwyl MacDonald for 3D modeling, Dave Hemer (UC Davis) for skillful machining, Robb Nichols (Aquarian Audio & Scientific) for advice and early hydrophone access, and Gary Elko and Jens Meyer (MH Acoustics) for advice on in-air ambisonics. They also thank Kelly Finn for surface observations and videography and insights on the appropriateness of contemporary animal-behavior observation protocols for revealing marine mammal behavior.

## Author contributions

J.P.C. designed and built the HAP underwater hydrophone recording array and underwater acoustic broadcast system with the expert advice of D.D.D. J.P.C. and D.D.D. conducted the field recording experiments and calibration of the recording and playback subsystems. A.M.J. was responsible for field photography and observations of surface behavior. These efforts were conducted using personal and University of California equipment. J.P.C., D.D.D., and A.M.J contributed to data acquisition, analysis, and interpretation. They also contributed to all aspects of manuscript writing and production.

## Data and materials availability

Data provided by the first author upon reasonable request.

Harpex is a trademark of Harpex Audio GMBH.

## Appendix A Ocean Bioacoustics

Sound propagation in water differs markedly and in key ways from propagation in air. Given human’s innate sense and experience of sound in air, the differences need to be taken into account when interpreting the signals that hydrophones pick up.

First, the speed of sound in water is five times that in air: 1, 500 meters per second compared to 340 meters per second, respectively, owing to the water medium being markedly denser than air. Practically, this leads to, for example, echos as sounds bounce off the seabed. Since density increases with depth, water depth is important and, of course, changes when changing anchorages. This also means that sounds from distant sources can be detected. For example, one is often surprised by the degree to which vessel noise is heard and in some cases dominates the undersea soundscape, even if vessels are not in sight. Commercial cruise liners are notable contributors to ocean noise given their immense displacement (key to waves generated by their passing) and massive engines.

Second, the precise nature of propagation in water is complicated by the fact that sound velocity increases with water pressure (and so depth) and decreases with water temperature and salinity.

Third, unlike sound in air, underwater sound at different frequencies propagates at different speeds—this is referred to as frequency dispersion. Thus, a distinct sound pulse detected at some distance loses its sharpness and blurs out over a time period much longer than the original pulse.

Taken altogether, the effects of these dependencies have on propagation are unlike those of our experience of sound in air. They often result in unusual and counterintuitive sound phenomena. The physics underlying these effects are nicely recounted in Ref. [10].

For example, the dependencies lead to a fascinating phenomenon of extremely long-ranged detection of sound signals in the ocean. This is the *Sofar channel*. Due to the competing effects of pressure and temperature on sound speed, there is a horizontal “channel” that conducts sounds like a waveguide: signals within a certain frequency band bounce between a shallow “ceiling” (perhaps 10s of meters in depth) and a “floor” (100s meters or more in depth). The net result is that sound signals in the Sofar channel can propagate very long distances—easily tens of kilometers or, depending on conditions, to hundreds or thousands of kilometers.

One the one hand, these properties mean that undersea sound is markedly more complex to analyze, understand, and harness. These complications add challenges both to successful life undersea and to directly monitoring and interpreting sound in the ocean. On the other hand, these properties also mean there is additional information available in ocean acoustic waves—information that can be harnessed for environmental sensing and for communication.

Given the undersea is their environment and given their evolution over millions of years, marine animals, such as whales, have accounted for and take advantage of these ocean-acoustic properties. These features affect what they can perceive, how they generate sound underwater, and how they communicate and socialize. Undoubtedly, many aspects of their vocalizations are naturally adapted.

Finally, these properties affect the acoustic signals one records via hydrophones and so, too, how one interprets what one is hearing.

Properties that largely determine sound propagation speeds and wavelengths in air and water are given in Table I.

## Appendix B Digital Audio Recording

The HAP acoustic signals were captured by a multi-channel digital audio recorder (Zoom F8n Pro) to high capacity SD cards.

Each of the HAP’s four channels were sampled at 24-bit resolution at a rate of 44.1 kHz. After a day’s collection of recording sessions, the audio data was transferred to a laptop and also to an additional backup disk.

## Appendix C Data Acquisition and Recording System

The complete data acquisition and recording system is shown in Fig. 4.

## Appendix D Spatial Audio Signal Processing Chain

The following outlines, in the sequence used, the signal processing steps developed to spatialize multichannel hydrophone recordings of underwater sound sources, such as the humpback whales who are the focus here. Section II B described the multichannel ambisonic hydrophone (HAP) recording device that produced the 4-channel spatial audio signals—so-called Ambisonic A Format. Familiarity with the HAP device and general ambisonic processing theory [21] is assumed.

1. The raw Ambisonic A audio files were recorded as WAV Poly format with a [Zoom F8n Pro digital audio recorder](https://zoomcorp.com/en/us/field-recorders/field-recorders/f8n-pro/). These were then separated into four mono WAV files using the [Wave Agent (Sound Devices)](https://www.sounddevices.com/product/wave-agent-software/) software application.
2. Each mono channel was separately “cleaned” in [iZotope RX10](https://www.izotope.com/en/products/rx.html) software to remove intermittent clicks and interference noise generated by the operating systems of the host sea vessel—i.e., sonar pinging, water filtration pump, navigation telemetry, and the like) and amplitude adjusted with identical settings applied to each Ambisonic A channel file.
3. The cleaned mono files were subsequently recombined into WAV Poly format and loaded into a channel of the digital audio workstation (DAW) [Reaper](https://www.reaper.fm/). Within Reaper, the Sparta Array2sh VST plugin was used to convert the Ambisonic A files to Ambisonic B (Fuma) format while adjusting for the speed of sound in water (approximately five times faster than in air) and correcting for the specific transducer positions on the 12 inch diameter spherical housing of the HAP array. These adjustments were essential to account for the propagation of sound in the ocean environment and the relative arrival times of sound waves to the individual hydrophone positions as necessary to maintain the precise phase accuracy of the ambisonic encoding.
4. Once the audio data files were converted into Ambisonic B (Fuma) format, they could be encoded into the variety of binaural, surround, or speaker dome configurations that the ambisonic format facilitates. The [Harpex X](https://harpex.net/) VST plugin was used to perform this encoding function due to its flexible interface and direct support for a variety of binaural and surround configurations that can also be visually displayed in real-time as a useful two-dimensional mapping of three-dimensional space.
5. An additional useful feature of the Harpex X plugin is its support for creating “beamforming” adjustments to increase the directionality of the encoded ambisonic signals. Multiple instances of the Harpex X plugin can be activated simultaneously to create multiple combinations of beamformed signals. This can be used to increase the amplitude sensitivity of the overall system with regard to sounds occurring at extreme distances.
6. The VST plugin host Audiomulch was used to simultaneously run multiple instances of Harpex X and to transparently mix the correlated output signals for assignment to various surround configurations or binaural output. A standard Head Related Transfer Function (HRTF) was used for conversion to binaural audio. In this case it was the widely used HRTF derived from the [Neumann KU 100 Binaural Dummy Head microphone system](https://www.neumann.com/en-en/products/microphones/ku-100/). See Fig. 3.

**FIG. 3:**
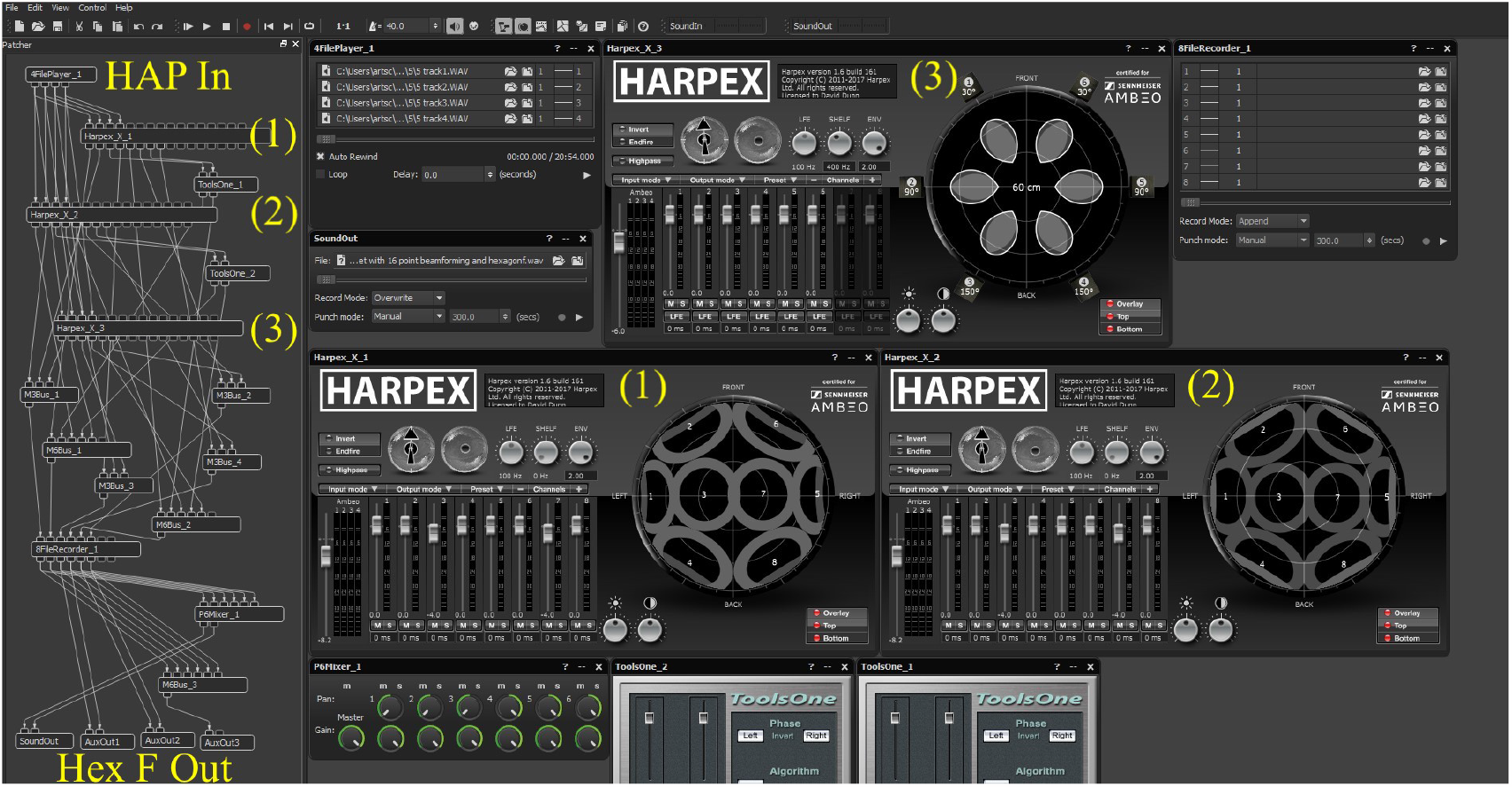
Spatial Audio Processing Chain: Multi-Harpex configuration within AudioMulch VST host for processing HAP first-order (4 channel) ambisonic A-Format hydrophone signals (HAP In) to Hexagonal F 6.1 surround format (3), using beamforming (1) and (2) to increase the directionality of the decoded Ambisonic signals and to steer away extraneous sound sources.

**FIG. 4:**
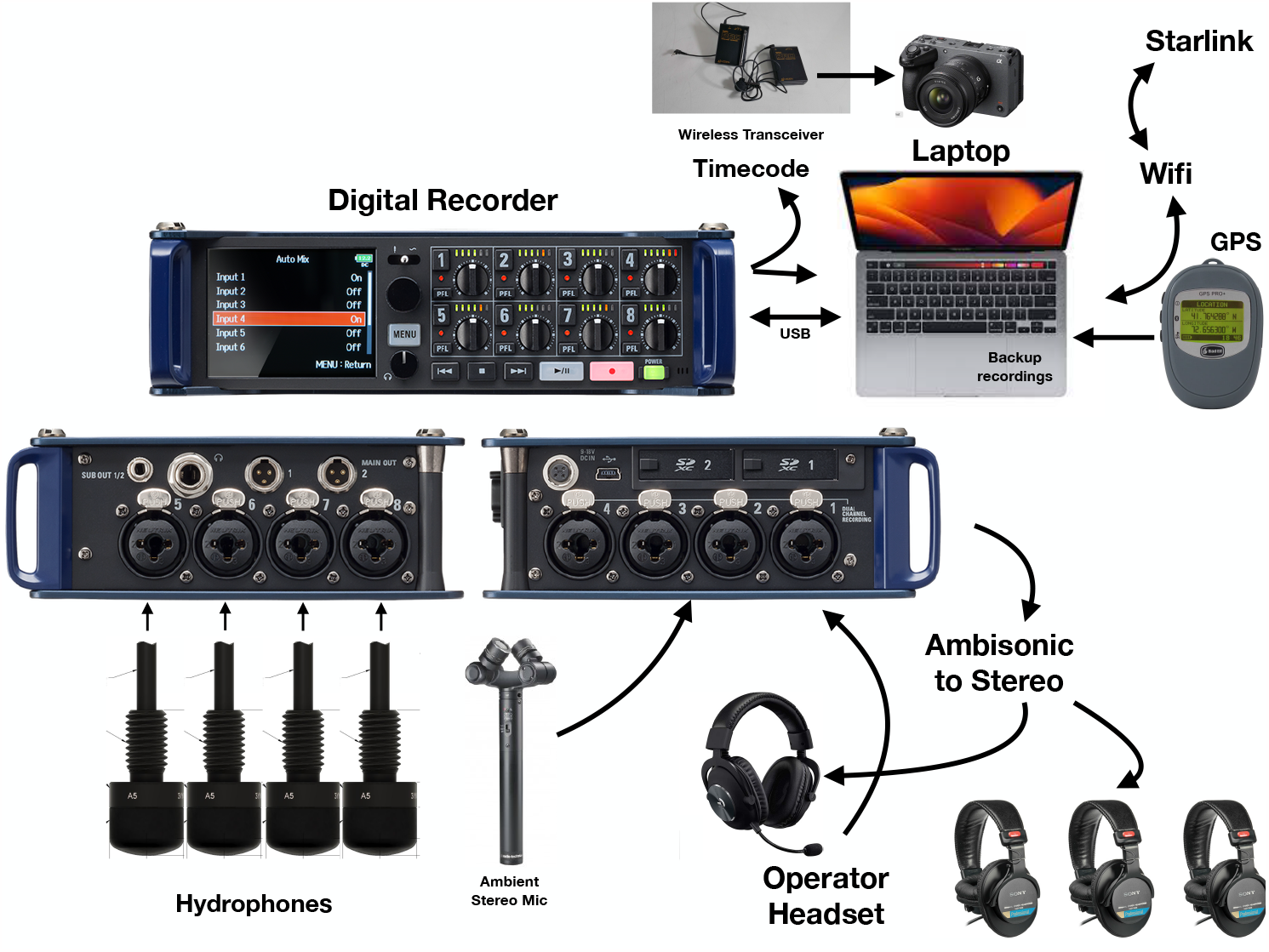
First-order Ambisonic Recording and Observation System.
7. All subsequent editing of the Ambisonic B or binaural audio files—prepared for public dissemination— was performed in either Reaper or [Adobe Audition](https://www.adobe.com/products/audition.html). No other equalization, filtering, signal processing (other than what has already been described), or juxtaposed mixing was used. For public presentation, cross-fading between selective event segments was employed to represent as much sonic and behavioral diversity as possible within a reasonable listening time frame.

## Appendix E Listening to Immersive Audio

We use headphones (Sony MDR-V6) to listen to the binaural renderings and a custom surround system to listen to the Hexagonal F 6.1 surround renderings. The latter consists of a subwoofer and six co-planar satellite speakers (located at head level and centered around the sweet spot where the listener sits).

Headphone listening is convenient and captures much of the spatial audio. However, Hexagonal F surround presents a richer sound environment with much improved directivity. The latter listening configuration is often necessary for discriminating sources in complex acoustic scenes and low-level or distant acoustic sources, as described in discussed in the humpback vocal harmonizing.

## Appendix F Surface observation

For completeness, we note that both video and photographic recording of surface behaviors were used simultaneously during the HAP recording sessions. The camcorder (Sony FX30 Digital Cinema Camera) and digital still camera (Nikon Z6 Mirrorless Camera, 400mm telephoto lens with teleconverter) were synchronized over a wireless link to the time-code generated by the digital audio recorder for the four HAP signal channels. Thus, surface observations and undersea acoustics could be accurately cross-referenced.

